# A comparative ‘omics’ approach for prediction of candidate *Strongyloides stercoralis* diagnostic coproantigens

**DOI:** 10.1101/2022.09.01.506149

**Authors:** Tegwen Marlais, Jack Bickford-Smith, Carlos Talavera-López, Hai Le, Fatima Chowdhury, Michael A. Miles

## Abstract

Human infection with the intestinal nematode *Strongyloides stercoralis* is persistent unless effectively treated, and potentially fatal in immunosuppressed individuals. Epidemiological data are lacking due to inadequate diagnosis. A rapid antigen detection test is a priority for population surveillance, validating cure after treatment, and for screening prior to immunosuppression. We analysed open access ‘omics’ data sets and used online predictors to identify *S. stercoralis* proteins that are likely to be present in infected stool, *Strongyloides-specific*, and antigenic. Transcriptomic data from gut and non-gut dwelling life cycle stages of *S. stercoralis* revealed 328 proteins that are differentially expressed. *Strongyloides ratti* proteomic data for excreted and secreted (E/S) proteins were matched to *S. stercoralis*, giving 1,057 orthologues. Five parasitism-associated protein families (SCP/TAPS, prolyl oligopeptidase, transthyretin-like, aspartic peptidase, acetylcholinesterase) were compared phylogenetically between *S. stercoralis* and outgroups, and proteins with least homology to the outgroups were selected. Proteins that overlapped between the transcriptomic and proteomic datasets were analysed by multiple sequence alignment, epitope prediction and 3D structure modelling to reveal *S. stercoralis* candidate peptide/protein coproantigens. We describe 22 candidates from seven genes, across all five protein families for further investigation as potential S*. stercoralis* diagnostic coproantigens, identified using open access data and freely-available protein analysis tools. This powerful approach can be applied to many parasitic infections with ‘omic’ data to accelerate development of specific diagnostic assays for laboratory or point-of-care field application.

**Author summary:** The worm *Strongyloides stercoralis* causes infectious disease in people throughout tropical and sub-tropical regions, leading to an extensive reduction in quality of life and even death. Millions of people are at risk of infection with this parasite and improved diagnostic and control methods and technologies are urgently required. Currently, most diagnosis is carried out through methods involving visual inspection of patient’s faeces, which has a number of drawbacks, particularly its poor sensitivity. This paper presents a new method to develop improved diagnostic tests for *S. stercoralis*, by computational analysis of publicly available gene and protein sequences to predict proteins that may be detectable in faeces. This would enable the development of rapid diagnostic tests in the form of lateral flows or dipsticks, with better predictive ability and fewer drawbacks than current diagnostic methods. A number of potential proteins, predicted to have all the desired characteristics for use in such tests were found through the new method and have been presented in this paper. With validation, new diagnostic tests for *S. stercoralis* could be developed from these results and the computational approach could be used to target other parasitic diseases.

## Introduction

The intestinal nematode *Strongyloides stercoralis* is a soil transmitted helminth (STH) prevalent in faecally-contaminated, humid soils in tropical and sub-tropical regions. Strongyloidiasis is estimated to affect up to 40% of people in many endemic regions [1, 2]. Infection occurs when infective third stage (L3) larvae penetrate the skin. The parasitic adult female resides in the epithelium of the duodenum where it feeds on host tissue. Although clinical signs may be mild or non-specific, long term infection by *Strongyloides* can have significant impact on quality of life and child development and progress to severe and fatal disease [3–5].

*S. stercoralis* is unusual among human parasitic nematodes in that it can complete its life cycle within the host, thus sustaining infection for decades if untreated [6–8]. During reduced immune competence due to immunosuppression, such as corticosteroid treatment of co-morbidities, or HTLV-1 co-infection, very large numbers of larvae may undergo this autoinfective cycle, causing hyperinfection or disseminated strongyloidiasis, both with a high fatality rate [9–11]. This autoinfective lifecycle means that infections will fully re-establish if worms are not completely cleared from the host during treatment. Diagnosis of strongyloidiasis and validation of cure after treatment are therefore imperative.

Treatment with the first-line drug ivermectin has a reported efficacy of between 57% and 100%. However, accurate determination of cure depends on follow-up and the diagnostic method used [8, 10, 12]. Albendazole and mebendazole, used to treat infection with other STH are less effective or ineffective against *Strongyloides* [13–15]. Moxidectin has shown effectiveness against *S. stercoralis*, that is equivalent to ivermectin, in early trials and continues to be developed [16, 17].

Diagnosis of strongyloidiasis can be made by microscopy using agar plate culture [18], Baermann funnel or spontaneous sedimentation, all of which isolate larvae from fresh stool [19, 20]. qPCR on extracted stool DNA is used in research and highly resourced laboratories, and improves sensitivity over microscopy [21–24]. However, sensitivity may be reduced by inadequate DNA extraction [25], low stool volume used in testing (as little as 0.1 g compared to 1 gram or more used for culture or larval concentration methods), and irregular larval excretion [12]. Serology, detecting antibodies to either whole worm or recombinant antigens NIE and SsIR, has sensitivity of 70-98% and high specificity [26] but can only distinguish cure months to years after effective treatment [27–29]. Therefore, there is a need for a rapid diagnostic test (RDT) that can be used for screening as well as for confirmation of cure. An antigen-based assay would fulfil this need, because it may achieve high sensitivity and specificity and test for active infection. Pilots of such assays have shown proof of principle under research conditions using antibodies against *Strongyloides ratti* and *Strongyloides venezuelensis* somatic or excretory/secretory (E/S) antigens [30–32]. However, identification of specific *S. stercoralis* protein antigens would enable production of standardised diagnostic tests on a large scale.

The wealth of ‘omic’ data now available in the public domain, coupled with online protein analysis tools, enables a computational approach to antigen discovery. This concept was termed reverse vaccinology when used in vaccine candidate discovery [33]. The approach begins with genomic analysis, as opposed to biochemical or serological methods and it also has significant potential in diagnostic antigen discovery. It has the advantages of not requiring culture of the organism and of revealing antigens that may be less abundant or difficult to purify *in vitro*. Incorporation of transcriptomic data can inform candidate gene/protein selection in parasites with multiple life stages, such as *S. stercoralis* [33].

Our approach was facilitated by the publication of the genomes of *S. stercoralis* and three related *Strongyloides* species [34]. Here, we have applied a series of computational analyses to open access transcriptomic, genomic and proteomic data from *Strongyloides* species and other helminths. We have used common bioinformatic tools, to identify *Strongyloides* protein antigens that may be diagnostic targets detectable in human stool, using a coproantigen capture RDT.

## Methods

### Data sources

This study used data obtained from public databases, primarily Wormbase ParaSite, a resource for parasitic worm genomics curated by the Sanger Institute and the EMBL European Bioinformatics Institute. Full details of data sources are listed in Supporting Information file S1.

*S. stercoralis* transcript sequences identified by the prefix ‘SSTP’ can be obtained via UniProtKB (www.uniprot.org) or WormBase ParaSite (WBPS: www.parasite.wormbase.org).

### Overview of method

Three criteria were applied for candidate antigen selection: presence in infected stool; specificity to *Strongyloides* and/or *S. stercoralis;* antigenicity, to facilitate raising sensitive antibodies. Datasets and computational analyses used to make this selection are detailed in Figure 1.

**Figure 1.**
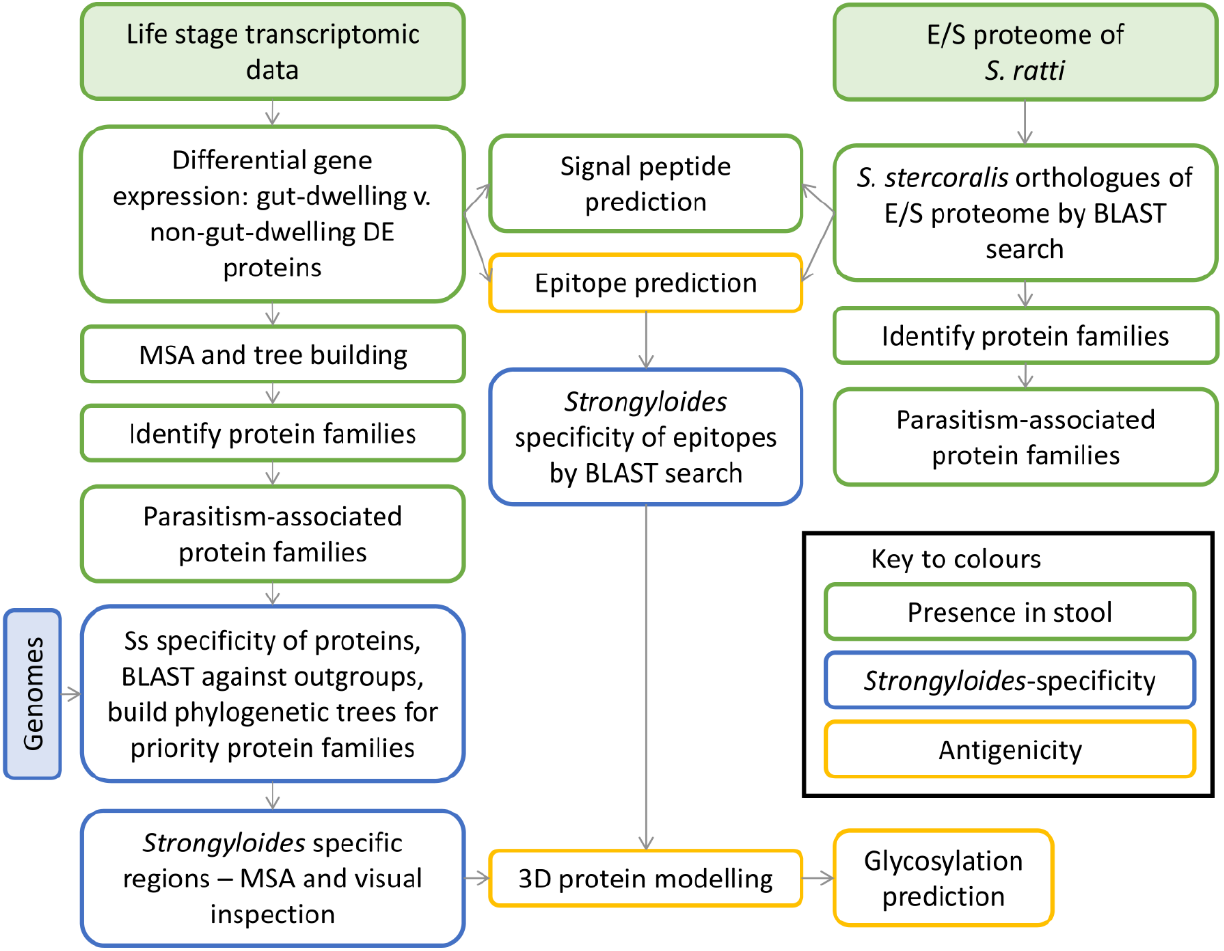
Schematic of the method for identifying coproantigens of *Strongyloides stercoralis* using ‘omic’ data and computational analyses. Filled shapes represent datasets, colours represent analyses against three characteristics of a candidate coproantigen. E/S, excreted and secreted; MSA, multiple sequence alignment; Ss, *Strongyloides stercoralis;* BLAST, basic local alignment search tool; DE, differentially expressed.

### Evidence for the presence of candidate coproantigens in stool

Analysis of the *Strongyloides* genomes by Hunt et al. (2016) [35] was used to identify protein families associated with parasitism. Five parasitism-associated protein families (‘priority protein families’) were then our focus for identifying coproantigens: sperm-coating-proteins/Tpx-1/Ag5/PR-1/Sc7 (SCP/TAPS), transthyretin-like (TTL), acetylcholinesterase (AChE), prolyl oligopeptidase (POP) and aspartic peptidases.

#### Transcriptomics

To reveal candidate coproantigens, we used transcriptomic data from Stoltzfus *et al*. (2012) [36], which analysed transcriptomes of 7 life stages of *S. stercoralis* (Figure 2). RNA data were downloaded from the National Centre for Biotechnology Information (NCBI) Sequence Read Archive (SRA), with triplicate reads from each stage. We grouped these data by life stage and by presence or absence within the host gut, thus representing stages excreting or secreting antigens into human stool (Figure 2).

**Figure 2.**
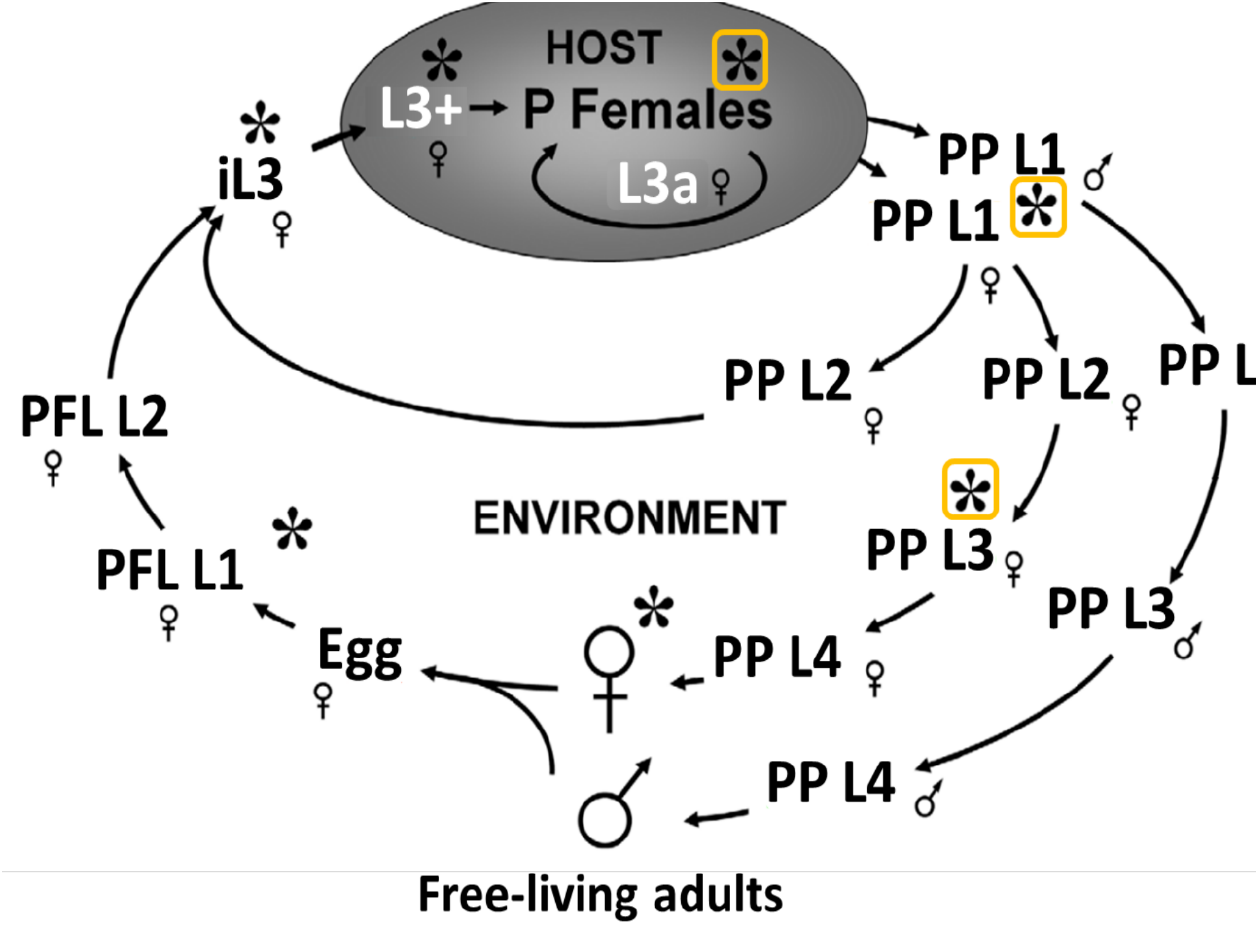
*S. stercoralis* life stages. Asterisks indicate life stages for which transcriptomic data were obtained by Stoltzfus *et al*. (2012). We grouped these transcriptomic data by presence in the host gut, (boxed asterisks) and outside the host gut. P Females, parasitic females; PP, post parasitic; FL, Free-living; PFL, post free-living; L1, stage 1 larva; L3, stage 3 larva; iL3, infectious third stage larva; L3+, tissue-migrating larva; L3a, autoinfective L3. Figure modified from Stoltzfus *et al*. (2012) [36] under a CC BY license. Accession numbers are given for NCBI SRA for the triplicate reads in Supporting Information S1 Table.

We calculated relative abundance of transcripts using RSEM [37] and bowtie2 [38] and subsequently separated RNA data from the 4 non-gut-dwelling stages and 3 gut-dwelling stages into two groups. Differential gene expression between the two groups was analysed using ebseq in RSEM. We selected genes that were differentially expressed with 100% confidence, in either direction, between gut-dwelling and non-gut-dwelling life stages.

#### Differentially expressed protein family identification

ClustalW [39] was used to perform multiple alignment of the differentially expressed (DE) genes and to produce a phylogenetic tree, which was annotated with iTOL [40]. The tree, labelled only with *S. stercoralis* gene accession numbers, facilitated grouping of the DE genes into protein/gene families but protein identities remained unknown at this stage.

DE proteins were firstly identified by submitting amino acid sequences of selected clusters on the tree to three domain-finding tools: Delta BLAST [41], InterPro [42] and ExPASy Prosite [43, 44] and a consensus of all three used to obtain probable protein identity. Secondly, DE protein sequences were submitted to BlastKOALA [45]. BlastKOALA protein identities were considered alongside previous consensus or used alone if there was no identity from three domain-finding tools.

#### Excretory/secretory proteomics

##### Source of proteomic data

Soblik *et al*. (2011) [46] submitted excretory/secretory (E/S) material of *S. ratti* parasitic females to mass spectrometry and identified the constituent proteins. In their presentation of the *Strongyloides ratti* genome, Hunt *et al*. (2016) [34] re-analysed the spectral data and obtained protein identities from corresponding genomic data of *S. ratti*. We acquired the list of parasitic female E/S proteins, with *S. ratti* genome accession numbers and protein identities, from Supplementary Table 19 of Hunt *et al*. (2016) [34] and subsequently obtained the corresponding amino acid sequences from the *S. ratti* protein file (WBPS v8) using samtools [47].

##### *S. stercoralis* orthologues to the *S. ratti* excretory/secretory proteome

At the time of this study, there were no E/S proteomic data available for *S. stercoralis*. Therefore, we obtained *S. stercoralis* orthologues of the *S. ratti* E/S proteins, by searching the *S. ratti* E/S proteins against a custom blast+ database consisting of the *S. stercoralis* protein file (WBPS v8), using blastp with word size 2 and e-value −50. *S. stercoralis* hits, in the form of accession numbers, were extracted from the resulting table and duplicated hits removed. Corresponding *S. stercoralis* amino acid sequences were extracted from the *S. stercoralis* protein file using samtools. VENNY 2.1 [48] was used to reveal the *S. stercoralis* accession numbers that occurred in both the DE proteins and the E/S orthologues. All the E/S orthologues were submitted to BlastKOALA as before, to obtain protein family identities, as well as matching them with the protein identities reported for the original *S. ratti* E/S proteins, by Hunt *et al*. (2016) [34]. Separately, differential gene expression data from analysis of the Stoltzfus *et al*. (2012) [36] dataset were extracted for the E/S orthologues that occurred in both datasets.

Signal peptide prediction for evidence that a protein is secreted was performed on the *S. stercoralis* DE proteins and E/S orthologues using SignalP 4.1 [49].

### Specificity of candidate coproantigens to Strongyloides

We used phylogenetic comparison to indicate *S. stercoralis* proteins with least homology to those of other relevant species, followed by multiple sequence alignment to identify exact regions of specificity.

A custom blast+ database was created from the genome-derived proteomes of selected outgroup species (Table 1). The outgroups were selected to represent parasitic and non-parasitic nematodes, as well as trematodes, cestodes and human. Human protein/coding sequence (CDS) data were provided by the Human Genome Project at the Wellcome Trust Sanger Institute and was obtained from Ensembl. Accession numbers for all data are available in Supporting Information file S1.

**Table 1.** Outgroup species used for phylogenetic comparison to *S. stercoralis*.

| Species | Rationale for selection |
| --- | --- |
| 1 *Ancylostoma duodenale* | Hookworm, prevalent nematode of humans |
| 2 *Ascaris lumbricoides* | Prevalent nematode of humans |
| 3 *Caenorhabditis elegans* | Free-living nematode and model organism |
| 4 *Clonorchis sinensis* | Trematode of humans |
| 5 *Enterobius vermicularis* | Prevalent nematode of humans |
| 6 *Homo sapiens* | Human, present in all samples |
| 7 *Necator americanus* | Hookworm, prevalent nematode of humans |
| 8 *Onchocerca volvulus* | Non-gut nematode of humans |
| 9 *Parastrongyloides trichosuri* | Relative of *S. stercoralis*, facultative parasite of opossums |
| 10 *Strongyloides ratti* | Close relative of *S. stercoralis*, serological cross-reactivity with *S. stercoralis* [50] |
| 11 *Strongyloides venezuelensis* | Close relative of *S. stercoralis* |
| 12 *Strongyloides papillosus* | Close relative of *S. stercoralis* |
| 13 *Syphacia muris* | Nematode of mice and rats, serological cross-reactivity with *Strongyloides* in experimental animal infection [51] |
| 14 *Taenia solium* | Cestode of humans |
| 15 *Trichuris trichiura* | Prevalent nematode of humans |

#### Specificity of differentially expressed proteins to S. stercoralis

Specificity of DE proteins to *Strongyloides* was assessed by their clustering in a phylogenetic tree after alignment with orthologues from the outgroups. The *S. stercoralis* DE proteins that resulted from the transcriptomic comparison were searched as separate protein families, against the custom outgroup database with blast+ criteria: word size 2 and e-value of −5 or −10, as appropriate, to obtain about 100 to 1,000 hits (S2 Table). In cases where there were very few DE proteins in a particular family, the DE protein(s) were also searched against a custom database consisting of only *S. stercoralis* genome-derived proteins (S2 Table). This was intended to increase the number of *S. stercoralis* proteins to enable species-specific clusters to be revealed on phylogenetic trees.

Protein hits from each outgroup, and the additional *S. stercoralis* hits where appropriate, were aligned with their respective original blast queries by multiple sequence alignment (MSA) using ClustalW, and phylogenetic trees were constructed. Trees were annotated with iTOL [40] to show proteins from each outgroup species in a different colour. *S. stercoralis* proteins which formed a distinct cluster, or clusters, on each phylogenetic tree were viewed in MSA along with the most and least similar proteins from each of the outgroups on that tree. These ClustalW alignments were analysed by eye for *Strongyloides* and *S. stercoralis-specific* regions which were submitted to BLASTP search against the NCBI nr, and Nematoda (taxid: 6231) databases, as relevant, to validate their specificity.

### Antigenic potential of candidate coproantigens

#### Epitope prediction

BepiPred 1.0 [52] and bcepred [53] were used to predict epitopes within the DE proteins and E/S proteome orthologues. A BepiPred threshold of 1.3 (range −4 to 4) was selected for maximum specificity of 96%, with corresponding 13% sensitivity, of predicted epitopes in order to minimise the chance of false positive predictions. Minimum length was 9 amino acids with no maximum. In the differentially expressed proteins, longer sequences with an overall very high epitope score were allowed to contain small regions scoring below 1.3.

Bcepred criteria were based on the reported highest accuracy of 58.7% which was achieved using a threshold of 2.38 for the average score of four amino acid properties: hydrophobicity, flexibility, polarity and exposed surface. In addition to BepiPred 1.0 and Bcepred, we also used BepiPred version 2.0 [54] for certain candidate antigens. This version became available only after the majority of the analysis and offered improved prediction of conformational epitopes. BepiPred 2.0 was used with the same epitope length criteria and an epitope score threshold of 0.55 (range 0 to 1) which provided specificity of 81.7% and sensitivity of 29.2% on epitope predictions.

Outputs from the two prediction tools were compared, initially for proteins present in both outputs. The predicted epitope regions of these proteins were then examined for sequence overlap. Prior to selection as candidate antigens, predicted epitopes were assessed for their specificity to *Strongyloides*. Sequences were searched using BLASTP against the NCBI non-redundant (nr) database. The “expect threshold” in BLASTP was increased if no results were obtained with default parameters. BLASTP output was examined by eye for the sequence identity and biological relevance, i.e. likelihood of presence in a human stool sample.

#### 3D modelling

Selected proteins of interest containing predicted epitopes, *Strongyloides-specific* regions, and in a priority protein family, were submitted to Phyre2 [55] for 3D structure modelling against known crystal structures, using the intensive mode. UCSF Chimera [56] was used to visualise and annotate 3D models to highlight specific sequences of interest on the model.

#### Glycosylation prediction

N-linked glycosylation was predicted with NetNGlyc [57] to account for the potential of a glycan to obscure protein antigen regions, or conversely to contribute to antigenicity. The prediction tool identified asparagine (N) residues with a high probability of being glycosylated via their amide nitrogen. Prediction was based on the motif N-X-S/T, where X is any residue except proline (P), and along with the presence of a signal peptide or trans-membrane domain on that protein, this indicates that potential glycosylation sites are likely to be glycosylated. Intracellular, intramembrane regions, or signal peptides of a protein are unlikely to be glycosylated. If present, a glycosylation site close to a candidate antigen region on the 3D protein could indicate that the protein is less likely to be accessible to antibodies in a capture assay and therefore a lower priority candidate, pending *in vitro* screening.

## Results

### Priority protein families

Hunt *et al*. (2016) [35] identified seven protein families associated with *Strongyloides* parasitism. For our coproantigen search, we focused on 5 of these, namely: sperm coating protein/Tpx-1/Ag5/PR-1/Sc7 (SCP/TAPS), transthyretin-like (TTL), acetylcholinesterase (AChE), prolyl oligopeptidase (POP) and aspartic peptidases.

### Differential gene expression in gut life stages

Of a total of 13,098 *S. stercoralis* genes identified in RNA-seq data by Stoltzfus et al. (2012) [36] we found 328 which we were 100% confident of differential expression between gut-dwelling and non-gut-dwelling life stages according to our groupings (Figure 3). Of these, 198 (60.4%) contained a signal peptide. Of the 328 DE genes, 203 were expressed more in gut-dwelling life stages than non-gut, including some or all of the proteins in the 5 priority protein families analysed here. These were therefore the focus of our coproantigen search (Figure 3 and S3 file). Twenty eight protein families were identified among the differentially expressed (DE) proteins, accounting for 193 (58.8%) of the proteins, with the remainder either not identified (22%) or given a disorder prediction (19.2%), indicating that they do not have a fixed conformation and are difficult to assign to a particular function or family (Figure 3).

**Figure 3.**
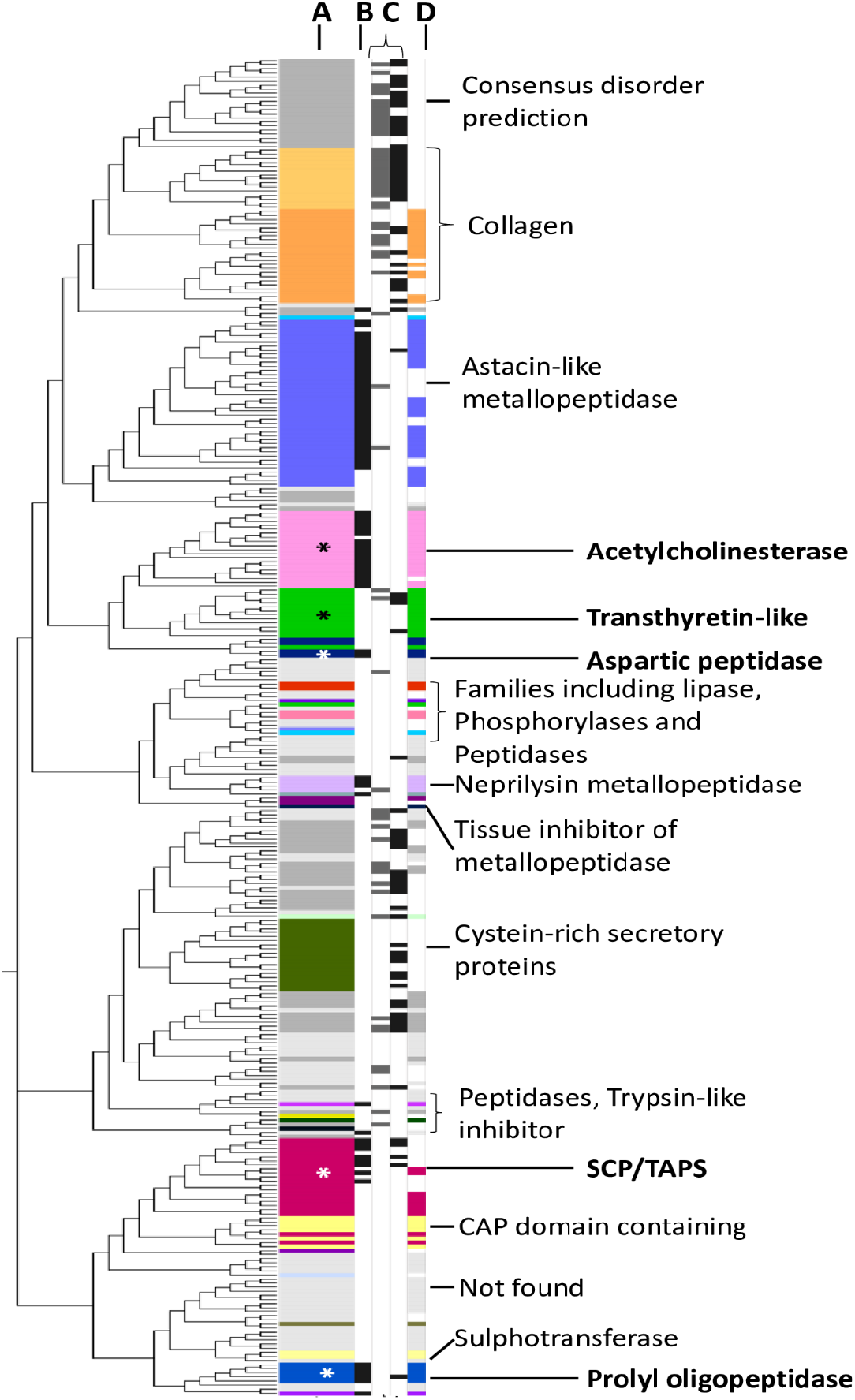
*S. stercoralis* proteins differentially expressed between gut-dwelling and non-gut-dwelling life stages. The total range of DE proteins is included for comparison. Proteins are grouped by similarity and families are identified. Any protein family with less than 2 BLAST hits present not labelled. Colours refer to features or families; the 5 priority protein families analysed (SCP/TAPS; TTL; AChE; aspartic peptidases, and POP) are labelled with an asterisk. Labels indicate layers representing: A, the 328 *S. stercoralis* DE proteins; B, proteins orthologous to *S. ratti* E/S proteins; C, proteins containing epitopes predicted by BcePred (left) and BepiPred (right); D, the 203 proteins that were DE in gut-dwelling life stages.

### Excretory/secretory proteome

In the absence of a *S. stercoralis* excretory/secretory (E/S) proteome, we identified 1,057 *S. stercoralis* proteins that had high homology to the 584 proteins in the published E/S proteome of *S. ratti* [34], of which 325 (30.7%) contained signal peptides. Original *S. ratti* E/S proteins were given as 582 accession numbers, however two were found to have alternative isoforms which are also included here. Multiple sequence alignment indicated that 550 (94.2%) of the 584 *S. ratti* E/S proteins had a *S. stercoralis* orthologue at the selected similarity level (e-value 1E-50) and that 284 (51.6%) of these had multiple homologues in *S. stercoralis* (S4 file).

To identify possible *S. stercoralis* E/S proteins among those differentially expressed in the host gut, we compared the 1,057 *S. stercoralis* orthologues with the 328 identified by transcriptomic data as DE in gut-dwelling life stages of *S. ratti*. Seventy seven (23.5%) proteins were shared between both data sets, of which 58 (28.6%) were shared between the 203 gut-stage DE proteins and the E/S orthologues.

To investigate gene expression of the E/S orthologues, we extracted data for the 1,057 *S. stercoralis* proteins from RSEM analysis of the entire 13,098 active *S. stercoralis* genes. Protein families were assigned by a combination of BlastKOALA, which identified 537 (50.8%) of the E/S orthologues, and the *S. ratti* protein identifications provided by Hunt *et al*. (2016) [34] (S5 file). Relevant proteins were assigned to the five ‘priority protein families’ (Figure 3).

### Predicted epitopes

Within the 328 confidently identified DE *S. strongyloides* proteins, BepiPred and bcepred jointly predicted epitopes in 104 proteins, 78 and 62 proteins respectively, with 36 proteins containing epitopes predicted by both tools (Figure 3 and S3 file). Within the 78 proteins, BepiPred predicted 125 epitopes, and within 62 proteins bcepred predicted 108 epitopes (S6 file). Fifty six epitopes contained overlap or identity between the two prediction tools (S6 file). Predicted epitopes ranged from 9 residues to entire proteins of up to 651 amino acids (aa) with BepiPred, and 8 to 66 aa with bcepred. These regions were given greater scrutiny in the context of species specificity and antibody accessibility.

*S. stercoralis* orthologues (n = 1,057) of the *S. ratti* E/S proteome were also submitted to BepiPred and bcepred which predicted 747 and 62 epitope regions respectively, in a total of 324 of the proteins. These ranged from 9 to 99 residues and contained 49 epitope sequences that overlapped, originating from 40 proteins (S6 file).

### *S. stercoralis-specific* candidates identified by phylogenetic comparisons

The five *S. stercoralis* protein families linked to parasitism and among the DE proteins were analysed for *S. stercoralis* genus or species-specificity. Separate BLAST searches of each protein family against 15 outgroups revealed the *S. stercoralis* proteins most likely to contain specific regions (Figure 4).

**Figure 4.**
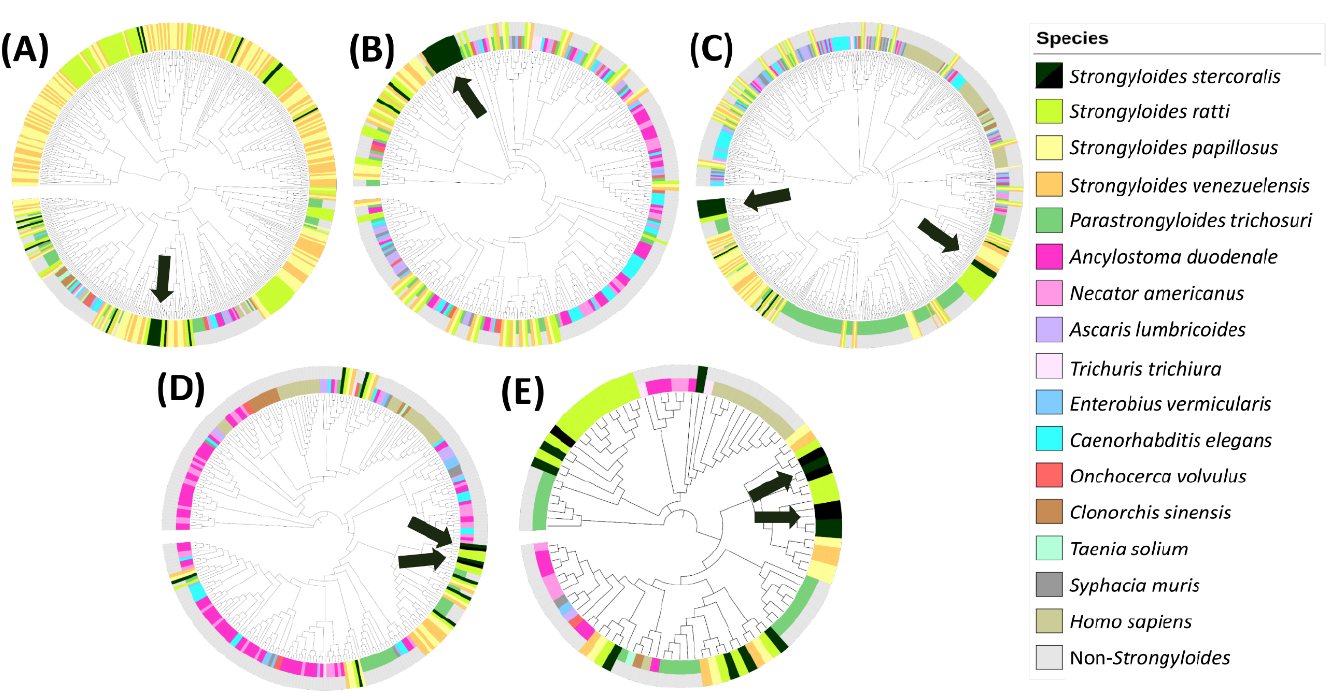
Phylogenetic alignment of *S*. stercoralis high priority family proteins with other *S. stercoralis* proteins and outgroups. Proteins of the five high priority families of *S. stercoralis* (dark green), from the differentially expressed dataset, in phylogenetic alignment with BLAST hits from *S. stercoralis* itself (black) and outgroups (other colours). Colours reaching the outer edge represent *Strongyloides* species, whereas shorter colour bands are *non-Strongyloides* outgroups. (A) SCP/TAPS, (B) transthyretin-like (TTL), (C) acetylcholinesterase (AChE), (D) aspartic peptidase, (E) prolyl oligopeptidase (POP). Arrows indicate clusters of *S. stercoralis* proteins containing species-specific regions; derived from both gut- and non-gut-dwelling differential expression data, augmenting detection of lack of species specificity.

*S. stercoralis* proteins from the clusters identified in the phylogenetic trees were examined in alignment with outgroup representative homologues with the most and least similarity. From the alignments, the *S. stercoralis* regions with least homology to outgroups were selected as candidate antigens. In some cases these included the entire protein. Results of this analysis are detailed below by protein family.

#### SCP/TAPS coproantigen candidates

Twenty one differentially expressed proteins were identified as definite (n = 19) or possible (n = 2) members of the SCP/TAPS protein family (Figure 3 and S3 file). The two ‘possible’ SCP/TAPS proteins clustered separately among CAP-domain proteins (Figure 3) and were therefore not included as definite members of this protein family. In phylogenetic analysis following alignment against outgroup genomes, seven of the 19 SCP/TAPS proteins formed a cluster of higher *S. stercoralis* specificity (Figure 4A, arrowed) (SSTP_0001008500, 8600, 8700, 8900, 0000511800, 512000, 513400). Multiple sequence alignment (MSA) and BLAST searching showed that these SCP/TAPS proteins contained several regions of apparent *S. stercoralis* species specificity, however, most were more highly expressed in non-gut life stages, such as tissue-migrating larvae. In total 8 of the 19 were expressed more in gut-dwelling life stages. Therefore SCP/TAPS from outside the cluster (in Figure 4A) were also viewed in MSA. A species-specific region was identified in gut-stage DE protein, SSTP_0000990000, consisting of a large 381 aa sequence of this protein which is upregulated in the parasitic female life stage (Table 2).

**Table 2.**
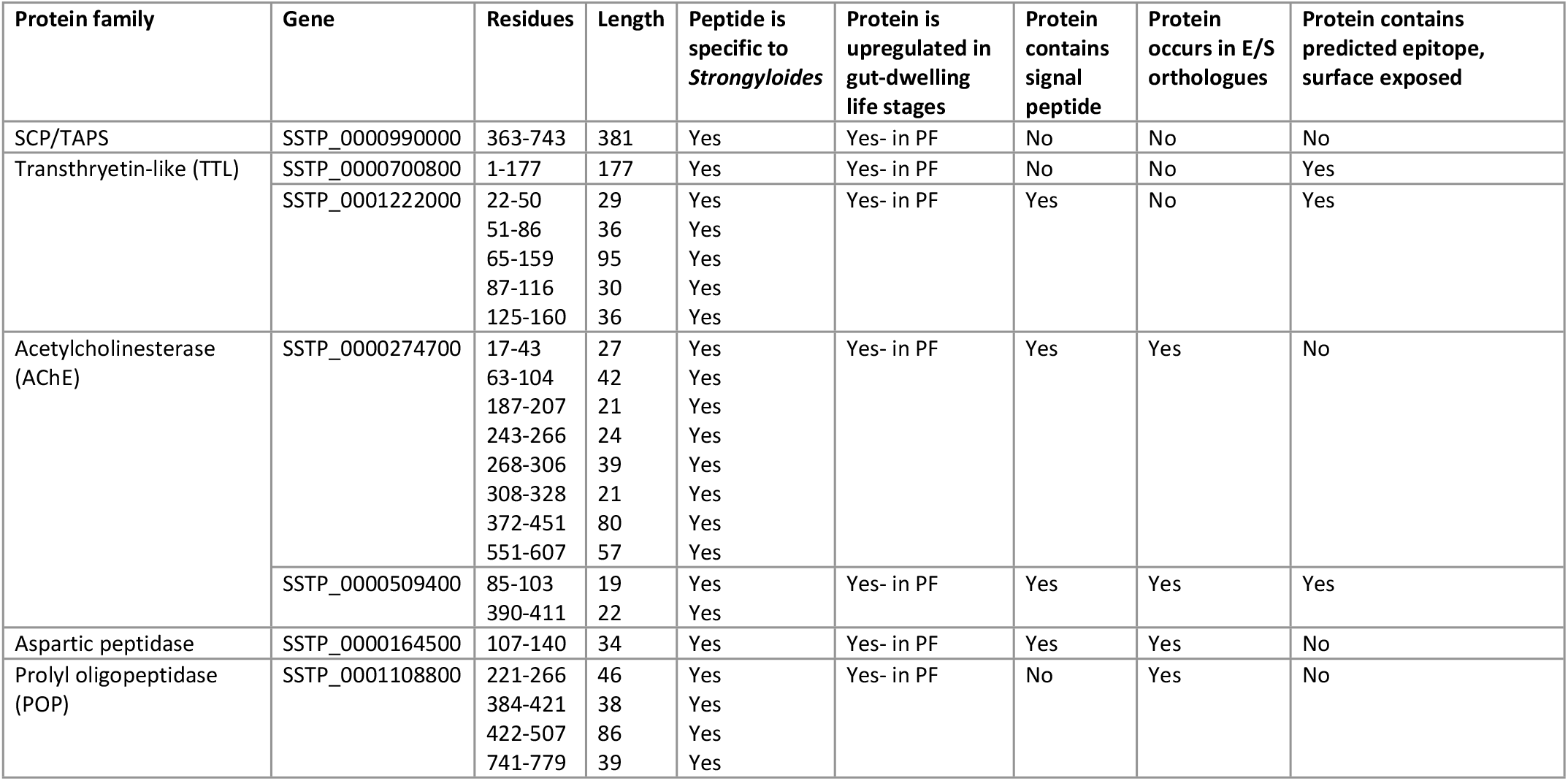
Candidate coproantigens of *S. stercoralis* based on upregulation in parasitism, amino acid sequence specificity to *Strongyloides* or *S. stercoralis*, and containing a predicted epitope. Gene accession numbers can be found in WormBase ParaSite or UniProtKB. Amino acid sequences are given in file S7. PF: parasitic female, E/S: excretory/secretory.

#### Transthyretin-like coproantigen candidates

Fourteen (14) TTL proteins were differentially expressed between gut and non-gut-dwelling life stages with 100% confidence, all of which were expressed more in gut-dwelling life stages, almost exclusively in parasitic female worms (Figure 3 and S3 file). Twelve of these proteins grouped as a cluster while the other 2 were more similar to other protein families or features (Figure 3). When aligned against homologous proteins from the outgroup species, 11 TTL proteins clustered together (Figure 4B, arrowed). All 14 TTL proteins were inspected visually in sequence alignment with selected outgroup proteins and five (SSTP_0000700800, 700900, 1222000, 0485800, 1133200) showed greater specificity to *S. stercoralis* and low sequence similarity to any of the outgroup homologues. However, when many of the possible *Strongyloides-specific* regions were BLAST searched separately, they were not sufficiently specific to be used as coproantigens. In particular, the search revealed that the amino acid sequence VTCDGKPL in protein SSTP_0000485800 is conserved across many nematode genera and should therefore be avoided in any candidate antigen. Two of the TTL proteins did contain one or more regions of *Strongyloides* species or genus specificity, giving a total of 6 TTL candidate coproantigens (Table 2).

In addition to high *Strongyloides-specificity* across its whole sequence of 177aa, TTL protein SSTP_0000700800 also contained predicted epitopes. To investigate the position of these, the protein was 3D modelled to a template generated from TTR-52 protein of *C. elegans* (PDB accession: 3UAF) (Figure 5). This model aligned to 89 residues (aa 2-90; 50% of the sequence) with 99.9% confidence. This protein contained a predicted glycosylation site at position N165 and, although not predicted to have a signal peptide, it was indicated as an ‘extracellular or secreted’ protein in UniProt [58] and is a known E/S protein family, therefore it is more likely to be glycosylated.

**Figure 5.**
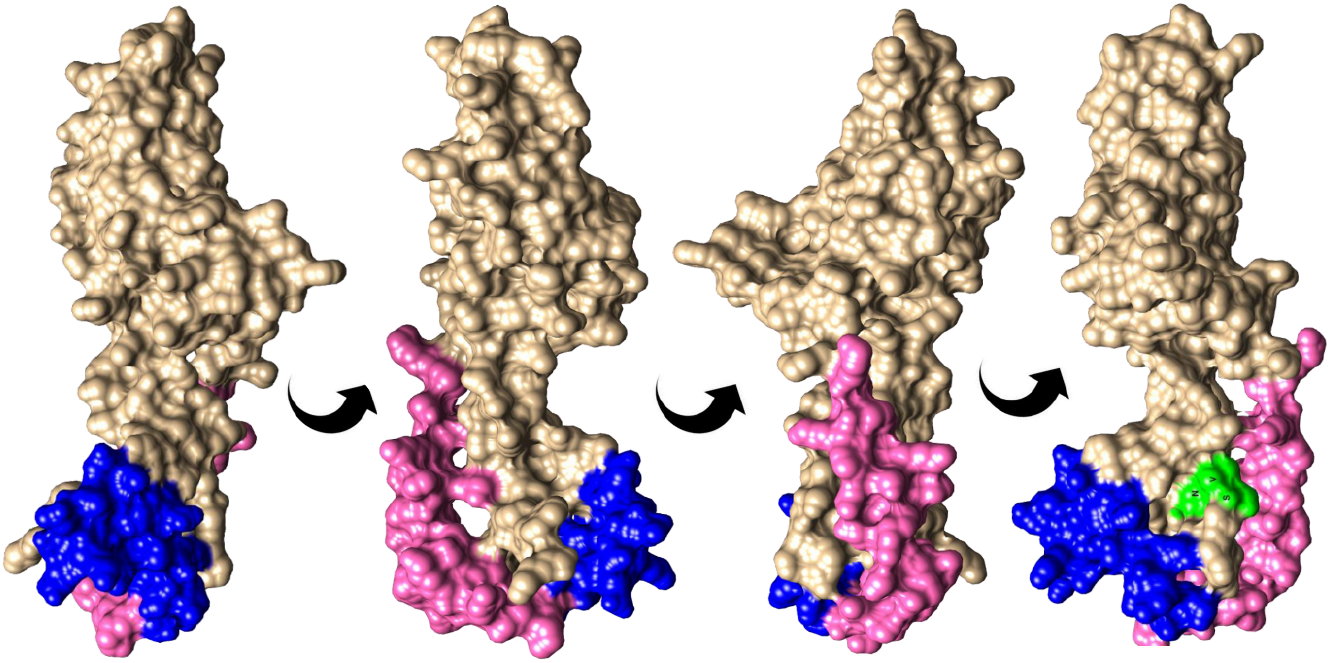
A transthyretin-like (TTL) protein of *S. stercoralis* in rotated view showing surface-exposed epitope regions and N-linked glycosylation sites. Accession number SSTP_0000700800. Blue, BepiPred predicted epitope aa’s 124-143; pink, bcepred predicted epitope aa’s 99-123; green, predicted N-linked glycosylation site at N165 in the motif NVS. The bcepred epitope (pink) extends to residue 141 but has been shortened to show the BepiPred epitope (blue), which overlaps with it.

#### Acetylcholinesterase (AChE) coproantigen candidates

Nineteen AChE proteins were in the DE dataset, of which 18 were expressed higher in gut-dwelling stages, particularly in parasitic female worms (Figure 3). The other AChE protein was highly expressed in infectious and tissue-migrating larvae (S3 file). All 19 proteins formed a cluster when compared with other DE protein families, indicating greater sequence similarity. The 19 proteins were then aligned with BLAST hits from outgroup species and most grouped into two distinct clusters in the corresponding phylogenetic tree, one cluster of 4 *S. stercoralis* proteins, the other of 10 (Figure 4C, arrowed), all of which were DE in gut-dwelling life stages. The cluster of 4 consisted of SSTP_0000274700, 671000, 638700 and 670800, all of which contained several regions of potential *S. stercoralis* specificity by visual inspection of the alignment. Equally, the 10 proteins within the other cluster contained multiple regions of possible *Strongyloides* specificity which were then analysed by BLAST to confirm this specificity. For AChE, 10 candidate antigens originated from 2 proteins: SSTP_0000274700 and 509400 (Table 2).

From the larger cluster in Figure 4C, one of the sources of candidate antigens, AChE protein SSTP_0000509400 was modelled with 100% confidence to 6 templates that jointly covered aa’s 17-551 (95%) with about 30% sequence identity. Peptides with potential *S. stercoralis* species-specific sequences were annotated on the model to view their surface exposure (Figure 6). BepiPred 1.0 and bcepred both failed to identify epitopes in this protein. However, BepiPred 2.0 did predict epitope regions, with moderate specificity and surface exposure. This AChE protein contained multiple potential glycosylation sites, three of which, at positions 38, 89 and 319, were predicted with high confidence on this known glycoprotein (Figure 6).

**Figure 6.**
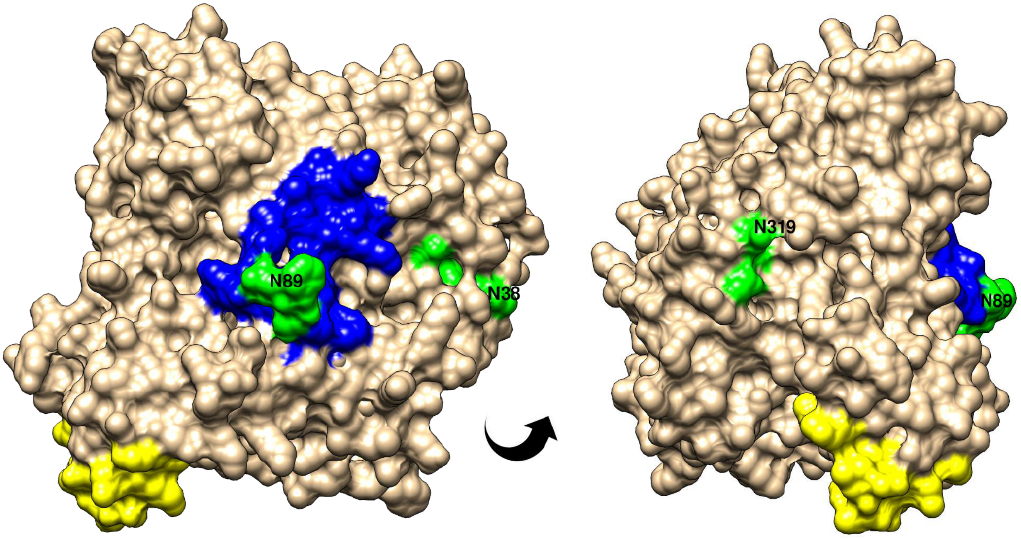
*S. stercoralis* acetylcholinesterase in rotated views showing surface-exposed epitope regions and N-linked glycosylation sites. Accession number SSTP_0000509400. Predicted features indicated in colour: blue, BepiPred 2.0 predicted epitope at residues 85-103; and yellow at 390-411; green, predicted N-linked glycosylation sites at N38 (NVT), N89 (NFS) and N319 (NLT). The site at N89 is within the blue epitope sequence.

#### Aspartic peptidase coproantigen candidates

Only 4 aspartic peptidases appeared in the DE proteins, all of which were expressed more in gut-dwelling life stages. Two of these were less reliably identified in this protein family so the other two (SSTP_0000164500 and 164700), with very high expression levels in parasitic female worms and both orthologues of E/S proteins, were analysed for *S. stercoralis* specificity. An additional 9 homologous proteins from *S. stercoralis* itself were included and all 11 proteins were then compared with the outgroups (Figure 4D, arrowed). The two DE proteins had low sequence similarity with each other. However, BLAST searching indicated that most of the possible species-specific regions lacked *Strongyloides* specificity and only one suitable peptide was identified from SSTP_0000164500 (Table 2). 3D modelling of this protein could not be achieved with any reliability and was thus not used for epitope selection. There were no predicted epitopes in either of the DE aspartic peptidases.

#### Prolyl oligopeptidase (POP) coproantigen candidates

The 5 differentially expressed POP proteins, all very highly expressed in parasitic females, were combined with a further 10 *S. stercoralis* homologues. In the phylogenetic tree, all 5 clustered with other *Strongyloides* proteins and away from *non-Strongyloides* outgroups (Figure 4E). Four of the DE proteins (SSTP_0000289100, 1108800, 1108500, 1019400) had higher species-specificity, clustering with other *S. stercoralis* homologues (Figure 4E, arrowed). One of these 4 proteins, SSTP_0000289100 contained two Bepipred epitopes and was very highly expressed in parasitic females, as well as being in the E/S orthologues (Figure 3 and S3 file). However, the epitope regions had moderate sequence identity to other nematodes, a *Staphylococcus* species and to *Plasmodium vivax*, suggesting widespread conservation. Therefore, these peptides were not considered sufficiently specific to *Strongyloides*.

Analysis of the POP proteins in multiple sequence alignment and subsequent search for similar proteins using BLASTP against NCBI nr and Nematoda databases, revealed several regions of high specificity to *S. stercoralis* in one of the DE proteins (SSTP_0001108800) (Table 2). Amino acid motifs conserved across genera, and therefore not *Strongyloides* specific, were removed from the sequences originally selected from the MSA, these included: DKLEN, KTDSK, RNAH and DIFAFI.

### Summary of candidate coproantigens

Table 2 presents selected candidate coproantigen protein regions that satisfy the criteria of being present in stool, specific to *Strongyloides* or *S. stercoralis*, and being antigenic. The full amino acid sequences of all candidate antigens are given in FASTA format in file S7.

## Discussion

The paucity of data on *S. stercoralis* infection prevalence and its low profile compared with the other STH species are largely due to inadequate diagnostics and lack of a single gold standard. While serology has the highest sensitivity for active disease, it is unsuitable for monitoring treatment outcome or defining cure in a timely manner [27, 59]. Incomplete cure and reinfection post-treatment may occur [8, 60, 61]. Therefore, we aimed to identify specific diagnostic targets from the nematode that could be captured by a rapid antigen detection test on stool samples. Such coproantigen assays are commercially available for *Giardia* and *Cryptosporidium* and have been developed for a wide range of human and animal parasites including *Ascaris* [62], *Fasciola* [63], *Echinococcus* [64], *Strongyloides ratti* [32], *S. venezuelensis* [30], *Opisthorcis* [65], *Toxocara* [66] and *Entamoeba histolytica* [67], among others. These assays employ either somatic, E/S material, or known antigens as targets.

We used open access data sources, published literature and freely-accessible online protein analysis tools to shortlist candidate antigens, based on the three criteria: presence in infected stool; *Strongyloides* or *S. stercoralis* specificity, and antigenicity. A similar study by Culma (2021) investigated cellular location, antigenicity and allergenicity, among other features, in the *S. stercoralis* proteome to identify potential vaccine and diagnostic targets [68]. Here, we focused on proteins that were differentially expressed between gut-dwelling and non-gut-dwelling life stages of *S. stercoralis*, according to RNA-seq data [36]. Seven protein families have been identified by other studies as expanded in the genomes of parasitic nematodes, and upregulated in parasitic life stages [35]. Studies of the *Ancylostoma* hookworm E/S proteome and *S. venezuelensis* somatic larval proteome detected some of the same families (SCP/TAPS, proteases and TTL) indicating likely presence in stool [69, 70]. To identify coproantigen diagnostic targets, we have performed a detailed analysis of 5 of the 7 parasitism-associated protein families. We also identified *S. stercoralis* orthologues of the *S. ratti* E/S proteome [34, 46].

We found an overlap of 77 proteins (5.9% of the total) between DE proteins and E/S orthologues. This limited overlap may reflect post-transcriptional control of expression [34]. Thus, grouping together transcriptomic data of life stages with different gene expression profiles may have limited our resolution of stage-specific coproantigens. E/S proteomics are an alternative starting point for coproantigen discovery. However, when we viewed gene expression data for all the E/S orthologues, no single life stage accounted for all the parasitic female E/S proteome. Although the *S. stercoralis* E/S orthologues broadly represented the *S. ratti* E/S proteome dataset, there were 1,057 compared to the original *S. ratti* 550 dataset. Therefore, differential gene expression between species is also likely to be a complicating factor and ultimately, the E/S proteome of the species of interest would be most suitable for coproantigen discovery. However, shared epitopes between these closely related species still warrant the use of all available data [50].

### Priority protein families and species specificity

Phylogenetic trees of *S. stercoralis* DE proteins and their homologues from a selection of outgroups focused our analysis on individual proteins from the priority protein families. These proteins were more likely to contain species- or genus-specific regions.

#### SCP/TAPS

The first of the priority protein families analysed here, SCP/TAPS, is among the CAP domain-containing proteins and is proposed to have a role in modulating the host immune response [35]. Here, we identified 21 SCP/TAPS proteins and 6 additional CAP-domain-containing proteins among the DE proteins. Many of the species-specific proteins or regions in this protein family, were highly expressed in tissue migrating larvae, rather than gut-dwelling life stages, therefore, only one protein expressed in adult female worms was the source of candidate antigens. This protein family has been studied in other parasitic nematodes, particularly the hookworms *Ancylostoma caninum, Ancylostoma ceylanicum* and *Necator americanus*, in which they are numerous. Multiple of these SCP/TAPS proteins have been demonstrated to be antigenic and recognised by anti-E/S antisera, showing that they do possess at least two of the characteristics predicted by this pathway.

#### Transthyretin-like proteins

Transthyretin-like proteins are a nematode-specific protein family and are known to be in E/S material of many species [71]. Two TTL proteins were sources of candidate coproantigens in the present study, both of which were expressed highly in the parasitic female worm and very little in other life stages. Neither of these DE proteins were among the E/S orthologues, however other TTL proteins were, and were expressed highly across multiple life stages. In addition, a TTL protein has been identified as a potential vaccine or diagnostic candidate through a similar bioinformatic pipeline, adding to the evidence for this protein family as a source of diagnostic targets [68]. The exact function of TTL proteins remains unknown in animal parasitic nematodes, but evidence from plant parasites suggests a role in subduing the immune response to favour worm survival [72].

#### Acetylcholinesterase

Secreted AChE has a possible role in enabling certain parasitic helminths to evade host expulsion mechanisms from mucosal surfaces [35, 73]. The transcriptomic and E/S proteomic data strongly supported this, with AChE family proteins in the E/S orthologues being expressed almost exclusively in the parasitic female life stage (S5 file). Secreted AChE differs from the neuromuscular protein in structure, gene family and substrate, being less specific to acetylcholine [73]. We found moderate homology between the predicted epitope regions of a *S. stercoralis* AChE and other nematode species. We identified candidate antigens in two AChE proteins. One of the candidate antigen proteins (SSTP_0000274700), contained 8 regions of *Strongyloides* specificity, when less specific regions were excluded, comprising over 50% of the full length 607 aa protein.

#### Aspartic peptidase

The aspartic peptidase family of enzymes is named for the creation of the active site from aspartic acid residues. Very few aspartic peptidases were DE, but had high expression in parasitic females and homologues to *S. ratti* E/S proteins. Aspartic peptidases play a role in haemoglobin digestion in other parasitic helminths, therefore secretion may be linked to feeding within the host gut epithelium where the adult worm resides [74, 75]. We identified a single candidate antigen from the few examined aspartic peptidases which had considerable homology to aspartic proteases from other *Strongyloides* species, but little to any other relevant species, whereas the full protein had high homology to multiple relevant nematode species including hookworm.

#### Prolyl oligopeptidase

Five POP family proteins were DE between gut and non-gut dwelling life stages. All 5 were very highly expressed in parasitic females where they may be involved in defending against the host immune and parasite expulsion response. In the trematode *Schistosoma mansoni*, POP enzymes have been found to cleave peptide hormones and neuropeptides [76]. Inhibiting POP activity in *S. ratti* lead to immobility in a concentration-dependent manner, even in *in vitro* conditions, indicating that this protein family is also vital to worm survival [46]. Only one of the 5 POP proteins contained regions of sufficient *Strongyloides* specificity for consideration as a coproantigen.

In addition to the shorter, specific sequences listed here, there is broader potential to express whole recombinant proteins such as those indicating higher *Strongyloides* specificity, in order to raise polyclonal antibodies for screening. Alternatively, specific monoclonal antibodies could be developed against these whole proteins and again, screened for *Strongyloides* specificity, as conducted by Abduhaleem et al. (2019) with a monoclonal antibody raised against *S. ratti* somatic antigens [77].

### Epitope prediction

We performed epitope prediction on the DE proteins and E/S proteome homologues using two open access online tools, which yielded many predicted epitope peptides. An alternative to this would be to scan the entire genome for epitopes, a method applied to vaccine candidate discovery [78]. The challenge faced by this approach is the complexity of conformational epitopes compared with linear peptide epitopes. Antibodies frequently bind to conformational epitopes formed by the 3D structure of the antigen, which therefore cannot easily be detected by sequence analysis alone [54].

The availability of 3D protein models can assist with selecting conformational epitopes by modelling a sequence onto the structure of a homologous protein and revealing adjacent amino acids on the surface of the protein [79]. Models do not necessarily have high sequence identity to the query sequence but this does not decrease the confidence in the model. Confidence in 3D models of >90% indicates that the protein adopts the overall folds of the model but may differ from the native protein in surface loops [55], thus this method is not guaranteed, but provides a good indication for selecting candidate antigens. *Ab initio*-modelled regions, where the sequence was not covered by the model, have very poor accuracy and should therefore be interpreted with caution and not used as the sole basis for selecting conformational epitopes. The field of 3D structure modelling has been progressed in leaps by the use of machine learning [80, 81].

We saw differences in predicted epitopes between the computational versus ‘by eye’ approach to selecting epitope regions. The DE protein dataset contained longer predicted epitopes due to the decision, where relevant, to extend predicted epitopes across a short region of lower epitope score whereas the computational selection worked only on the exact score threshold and would not join two adjacent high scoring regions.

Glycoprotein antigens were not considered in this study, apart from the presence of potential N-linked glycosylation sites on candidate protein antigens. Glycans form existing species-specific, highly antigenic diagnostic antigens, including CCA and CAA of *Schistosoma mansoni* and *Schistosoma* genus trematodes respectively [82], and LAM of *Mycobacterium tuberculosis* [83]. They have also been implicated in lysate seroantigen of *S. stercoralis* [84]. In helminths, glycan structures may not only be species-specific, but also life-stage specific [85]. Glycans may obscure some of the protein epitopes predicted here, particularly in the secreted AChE which is highly glycosylated. This is to be expected as glycans form many of the host-parasite interactions [85]. In addition, secreted candidate antigens may also contain O-linked glycans, via oxygen atoms of serine or threonine, which are not easily predicted. Although we have excluded potential glycan epitopes, they could be accounted for to some extent by expressing antigens of interest, ideally in a closely-related system, potentially *C. elegans* or even *Strongyloides* itself [86]. As an alternative, glycans could be excluded altogether by synthesising peptides or expressing recombinant proteins in bacteria, thus focusing the antigen search purely on proteins, as we have done here.

### Limitations and future work

We have described a methodological approach to discovery of diagnostic antigens. The process is predictive, based on computational analyses of ‘omic’ data. Predictions rely on the integrity of data sources, efficacy of software, and correct use of cut-off values at multiple stages of analysis. WormBase ParaSite, the source of most of the sequences used, is a curated database, regularly updated with high quality additional data and regarded as predominantly reliable. Thus, large-scale sequencing errors that might affect our use software, such as antigen prediction or BLAST analysis, are rare. Single nucleotide and small-scale sequencing errors are unlikely to affect results of the predictive pathways. With the E value cut-off used to select matching proteins by BLAST analysis, some proteins returned multiple hits. This may be due to paralogs or reflect an E value that is too lenient. Even in the latter case, over-inclusion of proteins at this stage, pared down by downstream analysis, is preferable to rejection of candidates. Proof of candidate antigen validity requires follow-up experimental laboratory research, which can reveal whether constraints and cut-off values have been too strict or too lenient.

Another limitation is prediction of antigens that are present in the stool. In our pipeline, a protein being excreted or secreted by a gut-dwelling stage of the parasite has been used as a proxy for presence in the stool. This rests on the assumption that the protein passes through the gut, is present in the stool in sufficient quantity and with antigenic properties unchanged by digestion or denaturation. As *S. stercoralis* establishes and matures in the small intestine, E/S proteins will not be subject to the denaturation and enzymatic degradation in the stomach but may be affected by enzymes in the small intestine. However, post-parasitic larvae, which hatch from eggs laid in the intestine, migrate through the small and large intestine, and can be found in faeces, so any E/S proteins produced by these stages, especially lower down in the intestinal tract, are less likely to be affected by host processes. The validity of these assumptions can be investigated by analysis of infected stool.

A further consideration is the focus of the approach on differentially expressed proteins and certain parasitism-associated protein families. There is rationale to targeting proteins with these characteristics, because they are proven to be expressed at higher levels in the relevant life cycle stages, are likely to be excreted or secreted and species or genus-specific due to adaptive radiation and protein family expansion. The focus on these groups limits the scope of an ‘omic’ approach, however, proteins that do not belong to one of these two groups might still have the characteristics of a candidate coproantigen. The approach presented here can therefore be applied to other helminth infections, but may be complimented by an expanded approach that uses whole genomic or proteomic data without focussing on parasitism-associated protein families or life stages, with the results of the two methods being used in conjunction.

Other potential proteins for coproantigen detection assays include: enolase, common to *Schistosoma japonicum* [87], *Echinostoma* and *Fasciola* [88], *Onchocerca* [89] and *Trichuris* [90], as well as in among *S. stercoralis* E/S orthologous proteins reported here and with constitutive high expression across all life stages (S5 file); protein 14-3-3 from E/S and somatic extract of *Strongyloides* [91, 92], *Ascaris* [93]*, Schistosoma* and *Ancylostoma* [70]. In addition, collagen, which forms some 80% of the outer cuticle of the nematode and was prominent among the DE proteins [94]. As an antigen, collagen would have to be carefully analysed for species specificity and the influence of glycosylation on its availability to capture antibodies [95]. A summary of E/S proteomic studies was presented by Ditgen *et al*. (2014) [96], and analysis of *S. venezuelensis* somatic proteins was made by Fonseca *et al*. (2019) [69] which may inform further antigen searches. Vaccine candidates for *S. stercoralis:* sodium potassium ATPase (SsEAT), tropomyosin, and a galectin (LEC-5) [97, 98] may also generate effective antibodies for antigen capture, if the relevant antigens are detectable in stool.

The most immediate work arising from this research is to validate the ‘omic’ pathway via wet laboratory analyses, specifically proteomic methods to detect and assess the antigenicity of proteins present in stool. Production of predicted candidates followed by antigenic testing could also be used to investigate the results of the ‘omic’ pathway as well as for development of lateral flow design for successful peptides. Results could be used to inform and refine results from the ‘omic’ pipeline, however, well-characterised stool sample collections are required to pursue diagnostic development.

Antigenic diversity must be considered, due to geographic differences between nematode strains [99]. This has impacted on diagnosis and vaccination for other parasitic infections [100, 101]. Diversity can be readily investigated by amplifying and sequencing the genes of interest from a wide geographic range of samples. For *S. stercoralis*, this is especially required, because the reference genome strain PV001 originates from a dog infection with UPD (University of Pennsylvania dog) strain [102].

### Conclusion

We have presented a detailed analysis of *S. stercoralis* proteins, leading to the selection of diagnostic coproantigens. We have identified multiple *S. stercoralis* candidate protein antigen sequences with evidence for their specificity to *Strongyloides* or *S. stercoralis* from phylogenetic and sequence comparison with relevant other species. Evidence supporting their presence in infected stool was assessed by belonging to parasitism-associated protein families, upregulation in gut-dwelling life stages, presence in E/S material of other helminths, and being among *S. stercoralis* orthologues of the *S. ratti* E/S proteome. Antigenicity was predicted using epitope prediction tools and 3D structure modelling. Peptides or whole proteins analysed and presented here form a selection of promising candidates for raising antibodies against and capturing *S. stercoralis* antigen in stool, with potential adaptation to prototype point-of-care rapid diagnostic tests.

## Author Contributions

Conceived and designed the study: MAM, TM, CTL. Carried out the study: TM, CTL, JB-S, HL, FC. Wrote the manuscript: TM. Reviewed and edited the manuscript: MAM, JB-S, CTL.

## Acknowledgements

We thank Dr Andrew Falconar for useful discussions and advice during this project.

Chimera is developed by the Resource for Biocomputing, Visualization, and Informatics at the University of California, San Francisco (supported by NIGMS P41-GM103311).

TM was funded by the Sir Halley Stewart Trust (http://www.sirhalleystewart.org.uk/). The views expressed within this report are those of the authors and not necessarily those of the Trust. TM was additionally supported by the John Henry Memorial Fund (UK Charity number: 1118007).

## Supporting information

**S1 Table. Data sources used.** CDS: coding sequence.

**S2 Table: Number of outgroup protein hits for S. stercoralis protein families from the differentially expressed (DE) dataset.**

**S3 file: *S. stercoralis* genes (n=328) differentially expressed in gut-dwelling (dark blue) versus non-gut-dwelling (light blue) life stages of the nematode with those containing predicted epitopes indicated.** Transcriptomic data from Stoltzfus *et al*. (2012), gut and non-gut groupings and differential analysis from the present study. Gene accession numbers (starting SSTP) are given for WormBase ParaSite or UniProtKB and life stage transcriptomic data accession numbers (starting ERR) for NCBI SRA. Expression levels are in normalized read counbts where red is high and white is low.

**S4 file: BLAST results: *S. stercoralis* orthologues of the *S. ratti* E/S proteome.** *S. ratti* E/S proteomic data from Hunt *et al*. (2016) was BLAST searched against the *S. stercoralis* genome-derived proteome using blast+ with e-value 1e-50

**S5 file: *S. stercoralis* orthologues of *S. ratti* E/S proteomic data.** *S. ratti* E/S proteins (from Hunt et al., 2016) were searched against a custom blast+ database consisting of the S. stercoralis genome-derived proteome (from WBPS v8) to obtain homologues.

E-value −50 and word size 2 were used to perform the search. Protein identities are given for the original *S. ratti* proteins (from Hunt et al., 2016) and the BLAST KOALA results from this study along with corresponding KEGG orthology. Gene expression in normalised read count is from analysis of data from different life stages of *S. stercoralis* (obtained from Stoltzfus *et al*., (2012)). Overlap is indicated with proteins which were also differentially expressed (DE) with 100% certainty between gut-dwelling and non-gut-dwelling life stages, as determined for this study. Proteins containing predicted epitopes according to two tools: BepiPred 1.0 and bcepred are indicated as well as proteins predicted by SignalP to contain a signal peptide. Accession numbers starting ERR refer to RNA-seq data in NCBI SRA and those starting SSTP are *S. stercoralis* gene transcripts which may be found in UniProt or WormBase ParaSite.

Life-stage abbreviations are explained in the manuscript text.

**S6 file: Predicted epitopes in *S. stercoralis* proteins using BepiPred and bcepred.** ES: Orthologues of

*S. ratti* E/S proteome. DE: Differentially expressed proteins between gut-dwelling and non-gut-dwelling life stages.

**S7 file: Candidate coproantigen sequences in fasta format.** FASTA headers are in the format: accession number, start aa, end aa.

